# Cooperativity of myosin II motors in the non-regulated and regulated thin filaments investigated with high-speed AFM

**DOI:** 10.1101/2022.02.24.481751

**Authors:** Oleg S. Matusovsky, Alf Mansson, Dilson E. Rassier

## Abstract

Skeletal myosins II are non-processive molecular motors, that work in ensembles to produce muscle contraction while binding to the actin filament. Although the molecular properties of myosin II are well known, there is still debate about the collective work of the motors: is there cooperativity between myosin motors while binding to the actin filaments? In this study, we used high-speed AFM to evaluate this issue. We observed that the initial binding of small arrays of myosin heads to the non-regulated actin filaments did not affect the cooperative probability of subsequent bindings to neighboring sites and did not lead to an increase in the fractional occupancy of the actin binding sites. These results suggest that myosin motors are independent force generators when connected in small arrays, and that the binding of one myosin does not alter the kinetics of other myosins. In contrast, the probability of binding of myosin heads to regulated thin filaments under activating conditions (at high Ca^2+^ concentration and with 2 μM ATP) was increased with the initial binding of one myosin, leading to a larger occupancy of neighboring available binding sites. The result suggests that myosin cooperativity is defined by the activation status of the thin filaments.

**eLife digest:** Muscle contraction is the result of large ensembles of the molecular motor myosin II working in coordination while attached to actin. Myosin II produces the power stroke, responsible for force generation. In this paper, we used High-Speed Atomic Force Microscopy (HS-AFM) to determine the potential cooperativity between myosin motors bound to non-regulated and regulated thin filaments. Based on the direct visualization of myosin-actin interaction, probability of myosin binding, and the myosin fractional occupancy of binding sites along non-regulated and regulated actin filaments, our results show no cooperative effects over ∼100 nm of the actin filament length. In contrast, there is myosin cooperativity within the activated thin filament, that induces a high affinity of myosin heads to the filaments. Our results support the independent behaviour of myosin heads while attached to actin filaments, but a cooperative behavior when attached to regulated thin filaments.

## Introduction

Myosin II is a non-processive molecular motor that binds to actin filaments to produce mechanical work, using the chemical free energy of adenosine triphosphate (ATP). After an initial attachment to actin, the myosin motor domain undergoes conformational changes associated with release of the ATP hydrolysis products inorganic phosphate (P_i_) and ADP from the active site of myosin. In this process, a force-generating power stroke, with swing of the myosin lever arm, is generated and there is a transition of myosin from the weak to the strong actin-binding states (Rayment et al., 1993; Fisher et al., 1995; Mansson et al., 2018; Robert-Paganin et al., 2020).

Myosin II molecules form bipolar filaments in skeletal, cardiac and smooth muscles and this filamentous form of myosin II allows the motors to collectively produce high forces during muscle contraction despite a low duty ratio (Finer et al., 1994; Ishijima et al., 1994; Yanagida & Ishijima, 1995; Kaya & Higuchi, 2010; Kaya et al., 2017; Pertici et al., 2018; Cheng et al., 2019; Cheng et al., 2020). The actin-attached fraction of the ATP turnover time, the duty ratio, is ∼ 5% (Howard, 1997), which enables high speeds of shortening (Pertici et al., 2018; Cheng et al., 2020). Although most studies looking to the mechanics of isolated myosin II have been performed with single molecules, assemblies of myosin II have been investigated in arrays developed with a small number of motors adsorbed to silica beads (Debold et al., 2005), optical fiber surfaces (Pertici et al., 2018) or with the native thick filaments (Cheng et al., 2020). These small ensemble studies show a load dependence and force-velocity relation that is similar to that observed in myofibrillar (Lowey et al., 2018) and cellular preparations (Edman & Hwang, 1977). Furthermore, these force-velocity relationships can be modelled using single molecule properties (Mansson et al., 2018; Mansson, 2019), and experimental data from single molecules (Kaya & Higuchi, 2010; Capitanio et al., 2012; Sung et al., 2015) suggesting that myosin II motors are independent force generators, as postulated decades ago (Huxley, 1957), even when they are attached to a common thick filament.

However, there are also suggestions that myosin molecules work cooperatively, and the work produced by motor assemblies is different from individual motors (Kaya et al., 2017). Accordingly, the attachment of one motor would interfere with the kinetics and attachment mechanics of other motors when working in arrays. The result casts doubt on the concept of independent force generators in motor assemblies. Cooperativity could also arise in double-headed molecules (Huxley & Tideswell, 1997; Brunello et al., 2007) or myosin motors that bind to adjacent actin sites (Caremani et al., 2013; Rahman et al., 2018). X-ray diffraction studies using muscle fibre preparations provide evidence that the coordinated movements of myosin heads may indeed regulate force generation (Irving et al., 1992; Linari et al., 2015). Finally, this form of cooperativity may arise from allosteric changes of the actin filament itself so that binding of one myosin molecule modifies the kinetics of myosin binding to nearby sites (Orlova et al., 1993; Tokuraku et al., 2009; Prochniewicz et al., 2010).

Other forms of cooperativity between myosin motors involve activation of the thin filament where several cooperative phenomena have been described (Gordon et al., 2000). In skeletal muscle sarcomeres, actin–myosin interactions are regulated by Ca^2+^ through the regulatory proteins troponin (Tn) and tropomyosin (Tm), that form the thin filament complex with actin. Each of the Tm molecules contact seven actin monomers and is associated with the three Tn subunits: Tn-T, Tn-I and Tn-C . Upon Ca^2+^ binding to Tn-C, conformational changes are triggered in the Tn–Tm complex resulting in a displacement of Tm that allows for myosin binding to actin (Galinska-Rakoczy et al., 2008; Lehman et al., 2009). We previously have shown that under relaxing conditions, thin filaments presented a combination of activated and non-activated segments along their lengths, and were not blocked from myosin; the equilibrium between blocked and closed states was defined by Ca^2+^-induced Tn-Tm conformational changes (Matusovsky et al., 2019). In addition, myosin binding to actin is also required for full activation, or to induce the open state of activation of the thin filament (McKillop and Geeves, 1993; Smith & Geeves, 2003; Desai et al., 2015). When myosin binds to actin, it may directly affect the regulatory system by changing the conformation of Tm, such that other myosin heads can attach to thin filaments (Geeves & Holmes; 1999; Gordon et al., 2000). Furthermore, the question remains if one or two myosin heads in a molecule are required for the full activation of the thin filament.

Therefore, cooperativity during myosin II-actin interactions can conceptually arise from at least two sources: cooperativity among myosin molecules within the thick filaments due to structural changes in the actin filament or cooperativity through activation of the regulated, thin filaments. Each cooperativity source may present different mechanisms. In this study, we used High-Speed Atomic Force Microscopy (HS-AFM) to evaluate the potential cooperativity of double-headed heavy meromyosin fragments (HMM) of myosin II that were connected through the S2 tail regions, while attaching between non-regulated actin filaments, or regulated thin filaments. Because HS-AFM allows the investigation of protein dynamics with nanometer spatial and millisecond temporal resolutions (Kodera et al., 2021; Heath & Scheuring, 2018; Matusovsky et al., 2021) our experimental approach allows us to investigate important aspects of myosin cooperativity, with a better resolution than previous fluorescence microscopy studies (spatial resolution limitation of >100 nm) (Desai et al., 2015) . Specifically for this study, we developed a method in which HMM motors, attached by their S2 regions to form a structure with up to 8-10 individual myosin heads (4-5 HMM molecules) bound to nearby sites along two actin filaments or two thin filaments (Fig. 1 and Fig. S1). The benefit of this approach is the ability to monitor the behavior of each of the HMM heads over the time of an experiment to evaluate the potential cooperative binding of HMM heads with either actin or thin filaments during the ATPase cycle. The approach also allows investigation of aspects of inter-head cooperativity as well as the potential to investigate cooperative changes along actin or thin filaments at spatial resolution similar to the inter-monomer distance along the filaments.

**Figure 1.**
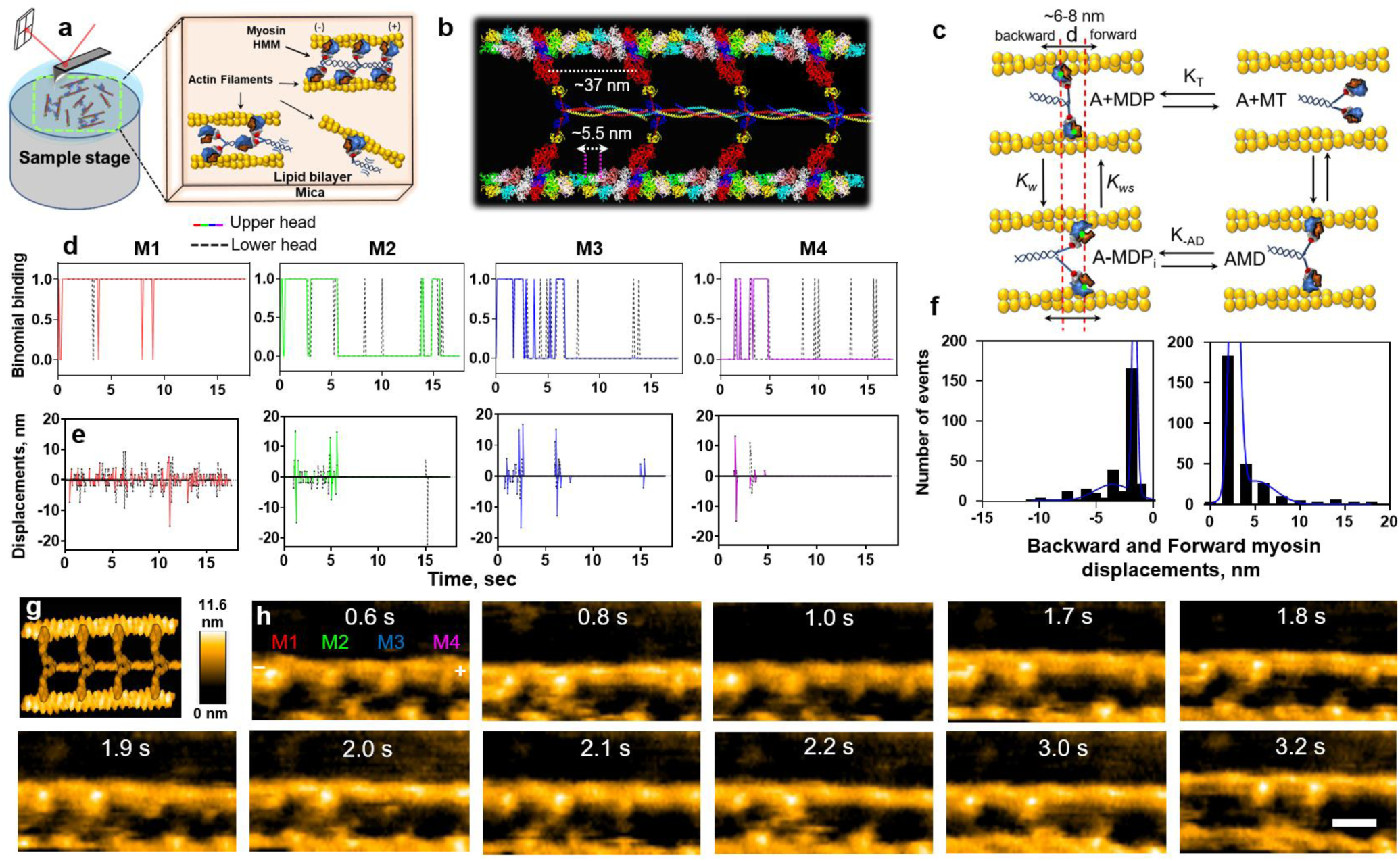
The kinetics of double-headed myosin motors bound to non-regulated actin filaments. (**a**) Diagram illustrating the approach used to study cooperative behavior of HMM molecules bound between two parallel filaments. (**b**) Structural model of actin-myosin complex for the experimental design used in the study; actin-myosin complex in rigor conditions (PDB:1M8Q) with upper and lower heads bound between two parallel actin filaments and attached by their S2 fragments (PDB: 2FXO). (**c**) Kinetics model of actin-myosin interaction and approach used to calculate the backward and forward myosin displacements; the green dots indicate the center of mass of the head. *K_T_* and *K_-AD_* are the constants of the ATP binding and ADP release; *K_w_* and *K_ws_* are the constants of the weak biding and weak-to-strong transition, respectively. (**d-e**) Representative time course of binomial binding (**d**) and head displacements (**e**) calculated for the individual HMM molecules (M1-M4) at the given time for upper and lower heads of each HMM molecule in the presence of 2 μM ATP. (**f**) The backward and forward myosin displacements in the F-actin-HMM complex (n=6, 589 events, ∼43 HMM molecules); data sets were fitted by sum of two Gaussians (r^2^=0.97 and r^2^=0.99, respectively). The two peaks for backward displacement: 1.8±0.2 nm and 3.7±2.0 nm; the two peaks for forward displacement: 2.7±0.5 nm and 5.1±2.1 nm. (**g**) Simulated HS-AFM image of the structural model shown in (**b**) performed in Bio-AFM viewer software (Amyot & Flechsig, 2020). (**h**) Representative HS-AFM snapshots of HMM molecules bound between two actin filaments at the indicated times (M1-M4 shown in color code duplicated in the **d-e** and across all of the figures), scale bar 30 nm. Related to Movies S1-S3.

## Results

### Experimental design to study myosin cooperativity by HS-AFM

In order to track the cooperativity behavior of the myosin heads within a sequence of successive HS-AFM images, we used an experimental approach in which pairs of HMM heads are attached to two actin filaments (Fig. 1, Fig. S1), as explained in details in a previous study from our laboratory (Matusovsky et al., 2021). Briefly, we aimed for an experimental situation in which two non-regulated actin filaments (F-actin) or two regulated cardiac thin filaments (cTFs) were bound to an underlying mica-supported lipid bilayer (SLB) surface, in parallel to each other, and with enough space for binding of double-headed HMM molecules between them. A cross-section analysis showed that the distance between two actin filaments during the experiments was 40.56 ± 9.65 nm, and the distance between cTFs was 67.69 ± 15.92 nm (Fig. S2). The observed difference in distance (27.1 ± 6.1 nm) between non-regulated F-actin and regulated cTFs did not affect the HMM binding and displacement analysis (Figs 1f, 2c).

**Figure 2.**
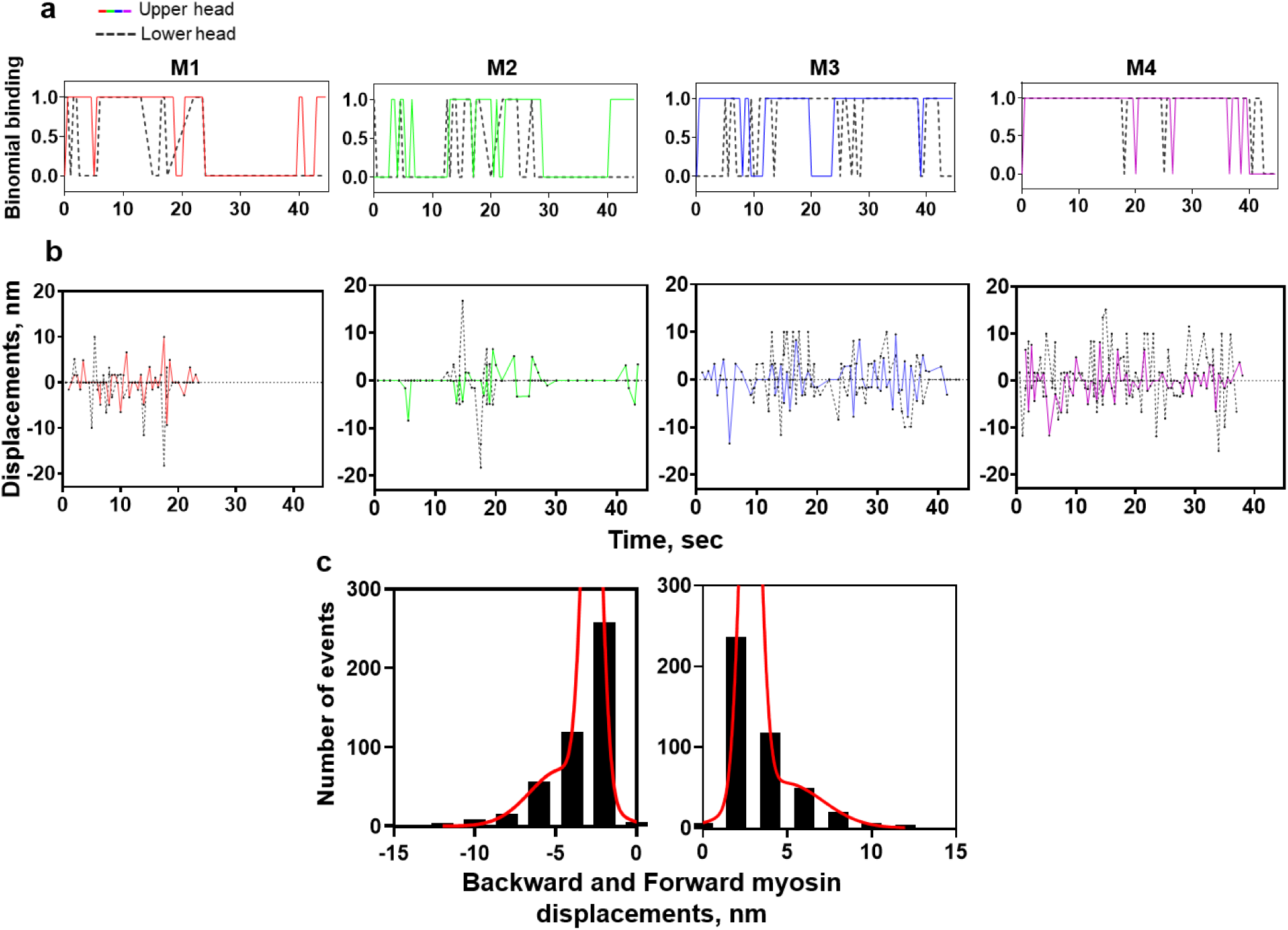
The kinetics of double-headed myosin motors bound to regulated cTFs. (**a-b**) Representative time course of binomial binding (**a**) and heads displacements (**b**) calculated for the individual HMM molecules (M1-M4) at the given time for upper and lower heads of each HMM molecule in the presence of 0.5 μM ATP and high Ca^2+^ concentrations. (**c**) The backward and forward myosin displacements in the cTFs-HMM complex (n=5, 911 events, ∼35 HMM molecules); data sets were fitted by sum of two Gaussians (r^2^=0.99 and r^2^=0.99, respectively).

Once the filaments were found in a parallel orientation, HMM fragments were added into the HS-AFM chamber filled with an experimental solution or placed on the top of the mica-SLB surface, in a solution containing 0.2-10 μM of NPE-caged or non-caged ATP. We then searched for events where each of the two HMM heads would interact with two parallel filaments. Immediately after both HMM heads were bound between parallel actin filaments, we activated the NPE-caged ATP in the solution by photolysis using a UV laser (340 nm) installed into the HS-AFM system (see Materials and Methods). The HS-AFM snapshots of two parallel non-regulated F-actins or regulated cTFs showed regularly bound HMM molecules between them in the absence or in the presence of ATP (Figs S1, S2). The globular upper and lower heads of each HMM molecule were bound in ∼30-37 nm proximity from each other, along the actin half-helical pitch structure (Fig. 1, Fig. S2). HMM heads were not bound to all the available actin-binding sites along the filaments at various experimental conditions, including rigor or in the presence of ADP in similarity to electron microscopy studies (Orlova et al., 1993). This observation may be related to the immobilization of S2 regions of each HMM molecule to the underlying lipid bilayer, allowing it to reach a maximum of ∼2-3 binding spots between neighboring actin monomers, i.e., 11 nm or 2 × 5.5 nm (Fig.1b). The ∼37 nm arrangement of myosin heads in our HS-AFM experiments is similar to the preferable binding sites of myosin heads along actin filaments (Steffen et al., 2001) and relate to the ∼37 nm hotspots for myosin head bindings along the thin filaments in the A-band of the sarcomere (Wang et al., 2021).

### Kinetics of actin-myosin interaction in the non-regulated and regulated systems

We characterized functional parameters of the myosin heads bound between two parallel non-regulated F-actins or regulated cTFs, including the average backward and forward displacements (*d*) of myosin heads in the presence of ATP (and high Ca^2+^ = pCa 4.5 in the case of cTFs). The HMM displacements calculated as a change in the center of mass (COM) of the myosin head at the given time during the experiment (Fig. 1c, see also Materials and Methods) were in the range of 6-8 nm. The backward (towards minus end of the filament) and forward (towards plus end of the filament) HMM displacements were calculated. The size distribution of HMM displacements revealed two distinct peaks in F-actin-HMM and cTFs-HMM complexes that most likely represent the events occurring through ADP (1-3 nm displacements) and P_i_ releases (over 3 nm displacements as previously described (Matusovsky et al., 2021). The sum of two peaks for backwards and forward displacements of HMM molecules were 5.5 ± 1.68 nm and 7.8 ± 1.96 nm on the non-regulated actin filaments, and 7.4 ± 1.73 nm and 7.6 ± 1.95 nm on the regulated cTFs (Figs. 1f and 2c).

The evaluation of displacement and working stroke of myosin heads in the HS-AFM was described in details in the Materials and Methods. Briefly, the working stroke is viewed as a transition from the weak to the strong-binding states evaluated by the changes in the lever arm movement. The change in the lever arm considers a defined polarity of the actin filament. In the presence of Mg.ATP, myosin heads detach from the filament and re-attach to the same or a new binding site, allowing us to determine the displacement of the myosin head by the change in COM. The calculated displacement in our study is slightly larger than the working stroke size of 5 nm reported for S1 (Capitanio et al., 2006) and slightly smaller than the values obtained from structural studies with single-headed myosin (∼10-12 nm) (Geeves et al., 2005). It is comparable with studies performed with myosin molecules evaluated with laser tweezers (Finer et al., 1994; Tyska et al., 1999) and single fiber mechanics (Piazzesi et al., 2002).

The representative binomial binding traces of the individual upper and lower myosin heads in the F-actin-HMM (Fig. 1d, Movies S1-S3) and in the cTFs-HMM (Fig. 2a, Movie S4) revealed that binding of one HMM head is not necessarily accompanied by the binding of the second HMM head for the given HMM molecule (M1-M4, Figs 1d and 2a). To specifically investigate the coordination between two heads in a molecule we analyzed the binding events at the different ATP concentrations. Tellingly, the binding events of two heads of given HMM molecule bound between two filaments was higher at lower ATP concentrations. At the higher ATP concentrations, the binding of either one head or two heads was approximately equally distributed (Fig. S3).

### Probability of HMM binding to the non-regulated and regulated actin filaments

To monitor the probability of binding events between individual myosin heads we applied a probability analysis based on a binary combination: HMM bound to the filaments equals to “1” and HMM detached from the filaments equals to “0”. To use this analysis, we need to evaluate if there is any directional bias in the myosin bindings along the F-actin and cTFs, either towards the barbed plus end or the pointed minus end of the filaments. Therefore, the polarity of F-actin and cTFs complexed with HMM was determined by using the morphology of the myosin heads bound to the filaments (Ngo et al., 2015). The bound myosin heads observed in the presence of Mg.ATP, Mg.ADP or in the rigor state allowed us to determine the polarity of the filaments (see Figs S1, S2-S5). According to our observations the most frequent myosin head orientation in the weak binding state (presence of Mg.ATP-γ-S) or strong binding state (rigor state) is the one where the heads of HMM molecules are positioned toward the minus end of the filament (Fig. 1h, Fig. 3a-b, Fig. 4b-d, Fig. S5). Therefore, binding events that occurred towards to the plus end of the filament (M_n_ ◊ M_n+1_) for individual upper and lower myosin heads at ATP concentrations ranging from 0.2-10 μM were used in the analysis.

**Figure 3.**
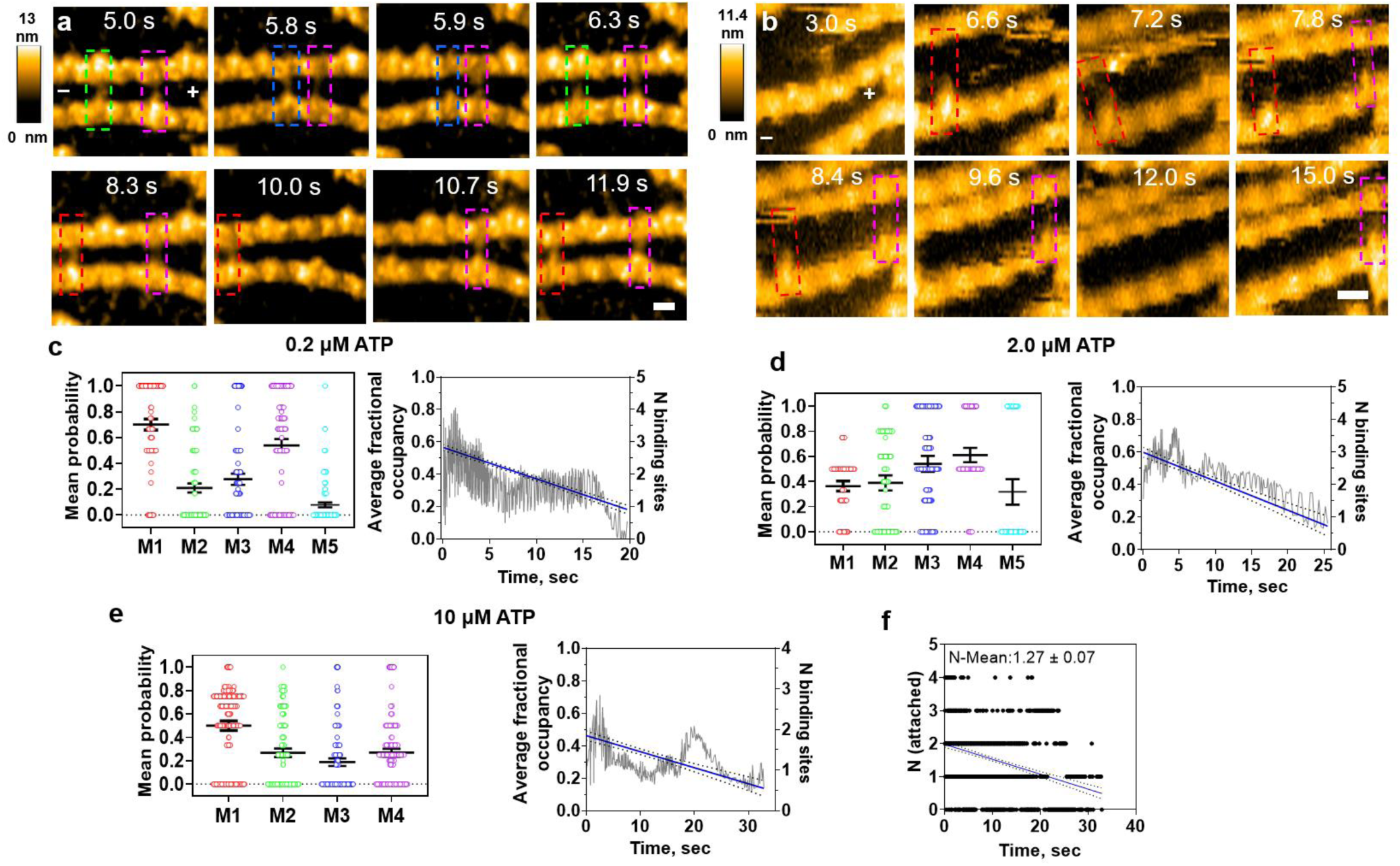
Probability of myosin heads binding to the non-regulated actin filaments. (**a-b**) Successive HS-AFM images of F-actin-HMM complexes, where HMM heads bind between two actin filaments in the presence of 0.2 μM ATP (**a**) or 2 μM ATP (**b**). Dashed color boxes indicate upper and / or lower HMM heads bound between two actin filaments. Numbers on each frame show the time in seconds. Related to Movies S1-S3. Scale bars: 30 nm. (**c-e**) Probabilities of the individual HMM head binding to the 4 or 5 binding sites on the non-regulated actin filaments in the presence of 0.2 μM ATP (**c**, 1235 events), 2 μM ATP (**d**, 567 events) and 10 μM ATP (**e**, 934 events). The average fractional occupancies by HMM heads for all of the sites for each of the ATP conditions showed a decrease in the occupancy with time (right panels in c-e, n=4). (**f**) Frequency distributions for the number of myosin heads attached to 4 neighboring binding sites along a given actin filament or thin filament over time (n=8 experiments, ∼35 HMM molecules, 898 events). Mean number (± 95% CI) of attached myosin heads relative to the 4 available binding sites given in the inset text. Full lines and dashed lines in the right panels in (**c-f**) represent the regression lines and 95% confidence intervals, suggesting a decline in the number of myosin molecules in time.

**Figure 4.**
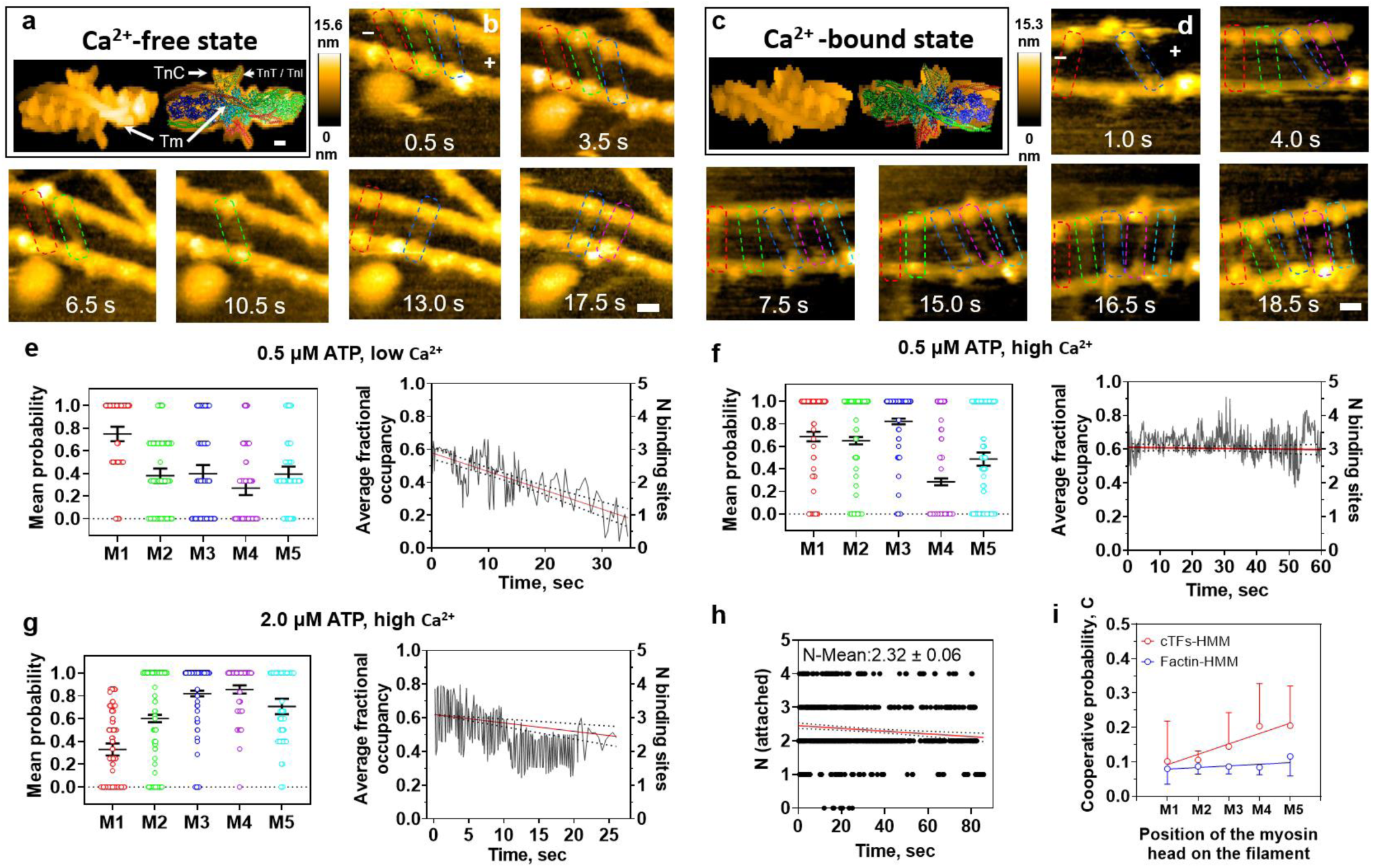
Probability of myosin heads binding to the regulated cTFs. (**a, c**) Fitting the simulated HS-AFM images (Amyot & Flechsig, 2020) of the cTFs at Ca^2+^- free (**a**) and Ca^2+^-bound (**c**) states into the cTFs molecular structures, scale bar: 2.5 nm. PDB: 6KN7 and 6KN8 for Ca^2+^ free and high Ca^2+^ states, respectively. (**b, d**) Representative HS-AFM snapshots of HMM molecules bound between two cTFs in the presence of 0.5 μM ATP and either low Ca^2+^ (**b**) or high Ca^2+^ (**d**) concentrations at the indicated times. Scale bars: 30 nm. Note, that the number of bound heads increase with time in (**d**). (**e-g**) Probabilities of the HMM heads binding to the regulated cTFs in the presence of 0.5 μM ATP and low Ca^2+^ (492 events) (**e**) or 0.5 μM ATP and high Ca^2+^ concentrations (2830 events) (**f**) or 2 μM ATP and high Ca^2+^ concentration (1839 events) (**g**) with the corresponding average fractional occupancy vs time shown in the right panels. (**h**) Frequency distributions for the number of myosin heads attached to 4 neighboring binding sites along all studied thin filaments over time in the activating conditions (n=7 experiments, ∼35 HMM molecules, 932 events). Mean number (± 95% CI) of attached myosin heads relative to the 4 available binding sites given in the inset text. Full lines and dashed lines in the right panels in (**e-h**) represent regression lines and 95% confidence intervals. The analysis suggested a decline in the number of myosin molecules with time, but to a lower degree compared to the F-actin-HMM complex. (**i**) Cooperative probability of binding of myosin heads in F-actin-HMM and cTFs-HMM complexes (n=7: cTFs-HMM n=8: F-actin-HMM, ∼40-50 HMM molecules in each data sets were analyzed; the data points shown as mean value ± 95% CI).

Initially, we tested binding of HMM between two filaments in rigor conditions, i.e., in the absence of ATP and Ca^2+^, or in the presence of ATP-γ-S, a slowly hydrolyzed analog of ATP (Fig. S5). At these conditions the HMM heads were tightly bound between two filaments with high fractional occupancies: ∼95% for the non-regulated F-actin and ∼79% for the native cTFs. The latter observation is consistent with the idea that the binding sites on cTFs in the absence of Ca^2+^ and ATP are present in an equilibrium between the blocked, closed and open states (Movie S4) (Matusovsky et al., 2019; Risi et al., 2017; Risi et al., 2021).

The probability analyses revealed that binding of M_n_ myosin head to the non-regulated actin filaments did not affect the subsequent bindings of the next M_n+1_ molecule (towards the plus end of the filament) in the presence of different ATP concentrations (Fig. 3, Movies S1-S3). While we can observe some random increase in the binding probabilities with 0.2 μM ATP or 2 μM ATP concentrations (Fig.3c-d) towards the plus end of the filament, the average fractional occupancy indicates a constant decrease in the occupancy of the binding sites with time, in all ATP concentrations used in this study (Fig. 3c-e, right panels). These data are consistent with the results pooled from 8 different experiments, suggesting that in the F-actin- HMM complex the most frequently observed events represent occupation of one binding site or no binding with the average number of occupied sites calculated as 1.27 ± 0.07 (Fig. 3f). These results suggest that HMM molecules are frequently detached from actin in the presence of ATP due to a lower affinity of myosin to actin in comparison to the affinity to the thin filaments (Fig. S4). This idea is consistent with the different time evolutions of the number of HMM molecules with actin and thin filaments (Figs. 3f and 4h) in the presence of ATP. It is also consistent with findings that the fractional occupancy of actin-binding sites in the absence of ATP (rigor) or presence of slowly hydrolyzed ATP-γ-S did not change with time (Fig. S5) when HMM is bound to both actin and the thin filaments all the time, suggesting that the myosin heads did not detach from the filaments over the time of the experiment due to interaction with the scanning cantilever tip.

At the non-activating conditions with thin filaments in the blocked state (0.5 µM ATP, pCa 9.0), myosin heads revealed a similar decrease in the binding probability and fractional occupancy (Fig. 4e) compared to the bare F-actin-HMM complex. In contrast, the probability of myosin heads binding to cTFs in the closed state under activating conditions (0.5 µM ATP, pCa 4.5) was increased (Fig. 4f-g, Movie S4) compared to myosin heads binding to actin filaments (Fig. 3c-d). This feature is reflected in the increased average number of attached myosin heads in the cTFs-HMM complex under activating conditions, almost doubling the average number of bound heads (Fig. 4h) compared to the situation with the F-actin-HMM complex (Fig. 3f). The increase in probability of binding of the myosin heads to the cTFs is also matched in the kymograph images of cTFs-HMM complex at the high [Ca^2+^] and different ATP concentrations, when compared to the kymograph images obtained from F-actin-HMM complex at the various ATP concentrations (Fig. S4). The random presence of activated and non-activated sites across cTFs at the relaxing (pCa > 8) or activating (pCa 4.0-4.5) conditions (Risi et al., 2017; Matusovsky et al., 2019) complicated the analysis and can explain the pattern of varied mean probabilities between HMM molecules (Fig. 4f-g).

### Cooperativity in the non-regulated and regulated actin-myosin systems

To quantify the observed probability binding pattern in the binary system, we applied the following equation 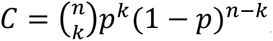, where C denotes cooperative probability of binding between neighboring myosin heads, n = number of total events or subsequent frames of the experiment; k = number of binding events in the subsequent frames, p = probability of binding, *i.e.* the ratio between binding events and total events, and 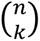 represents the combination of total and binding events expressed as 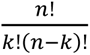 (see Supplementary Table 1).

Following this analysis, we found no change in the probability of cooperative binding of the HMM heads to the non-regulated actin filaments. It suggests that the interaction in the F-actin-HMM complex is largely random. The linear regression slope of the cooperative probability binding between neighboring myosin heads in the F-actin-HMM complex showed no significant deviation from zero (p = 0.295) with the Pearson’s r = 0.766 and r^2^ = 0.59. In contrast, the linear regression slope of the cooperative probability binding between neighboring myosin heads in the cTFs-HMM complex showed significant deviation from zero (p = 0.022) with the Pearson’s r = 0.953 and r^2^ = 0.91 (Fig. 4i). In accordance with these results, the individual fitting for each experiment demonstrated an increase in the cooperative probability of binding of the HMM heads to the regulated cTFs in comparison to that of in F-actin-HMM complex (Figs S6 and S7). This is broadly consistent with cooperativity, although the degree of cooperative binding in the cTFs-HMM was variable between experiments as can be noticed from confidence intervals (Fig. 4i).

## Discussion

In this study, we used a binomial probability analysis to evaluate the potential cooperativity between myosin motors while attached to actin or regulated thin filaments. Our experimental approach – using myosin motors that can attach between two filaments positioned in parallel on the surface - is particularly well-suited for this analysis, and we could visualize several different motors at the same time. Despite this approach has geometrical features that are distinct from those of the actin-myosin arrays operating within a muscle sarcomere, it has been recently shown that each of the double myosin heads can acquire different lever arm confirmations and bound two different thin filaments in rigor (Wang et al., 2021). This is also true when myosin attaches to actin and thin filaments in the presence of ATP, when each head may be in a different state, at a given time of the ATPase cycle (Matusovsky et al., 2021). This feature may enable myosin double heads to interact with two different thin filaments within the sarcomere, potentially maximizing muscle power and efficiency. Of particular interest, is the fact that myosin motors can be assumed as the independent force generators even when connected in small assemblies.

The displacements produced by each individual myosin head within a given HMM molecule in our experimental conditions were in the range of ∼6-8 nm, slightly larger than the working stroke size of 5 nm reported for myosin S1 (Capitanio et al., 2006) and slightly smaller than the values obtained from structural studies with single-headed myosin (∼10-12 nm) (Geeves et al., 2005). It is comparable with studies performed with myosin molecules evaluated with laser tweezers and single fiber mechanics (Finer et al., 1994; Tyska et al., 1999; Piazzesi et al., 2002). Therefore, our results are consistent with studies utilizing different techniques.

### Cooperativity between myosin motors

There are several studies suggesting that myosin molecules work cooperatively (Vilfan & Duke, 2003; Hilbert et al., 2013; Kaya et al., 2017; Hwang et al., 2021), and that the work produced by assemblies of motors is different from individual motors. For instance, a study using synthetic myosin filaments measured 4 nm stepwise actin displacements at a high load (>30 pN). Due to the fact that the mechanical work of 4 × 30 pN nm = 120 pN nm ≈ 30 k_B_T (k_B_: Bolzmann constant; T absolute temperature) is greater than the free energy of Mg.ATP turnover (25 k_B_T), the authors concluded that the steps they observed could not be produced by single motors but potentially due to coordinated force generation by several myosin motors (Kaya et al., 2017, Hwang et al., 2021). Despite the fact that theoretical analysis (Duke, 1999; Månsson, 2020) suggests that this finding is consistent with previous models of independent force generators as proposed previously (Huxley, 1957; Huxley, 1988; Hill, 1974), it casted some doubt on this concept when motors work in arrays. In this regard, the present study is consistent with fully independent force-generators along the actin filaments. Importantly, the HS-AFM approach allows us to demonstrate that this applies for neighboring actin target zones separated by ∼37 nm, appreciably shorter than possible to resolve under dynamic conditions using fluorescence microscopy (e.g. Desai et al, 2015). This result does not seem to be consistent with previous findings suggesting that binding of a myosin head allosterically affects the properties of the entire actin filament with potential changes of myosin affinity at other sites (Tokuraku et al., 2009). However, because we have only performed our studies under a limited number of specific conditions, we cannot completely exclude that such allosteric effects occur under certain conditions, e.g. at submicromolar concentrations of Mg.ATP as in some of the experiments of Tokuraku et al 2009. In contrast to the results with pure F-actin we found strong evidence for cooperativity between neighboring thin filament target zones as further considered in detail below.

Another form of cooperativity has been suggested by X-ray diffraction studies using muscle fibre preparations, indicating that the two heads of a given myosin molecule may bind sequentially to resist stretch of the active muscle (Brunello et al., 2007). Such sequential actions of the two heads have also been suggested (Edman et al., 1997, Huxley & Tideswell, 1997; Conibear & Geeves, 1998) to occur during shortening to account for rapid repriming of the myosin power-stroke after a quick release, high power output during shortening and other phenomena. To the best of our knowledge interhead cooperativity has, however, not previously been observed experimentally under dynamic conditions in the presence of ATP. Our demonstration that binding of one myosin head increases the probability for binding of the second head is thus unique by demonstrating the potential for inter-head cooperativity where binding of one head increases the probability of binding of the second head to another. The demonstration for this potential is of interest despite the fact that the distance between neighboring, roughly parallel actin filaments in our study is appreciably larger than in the muscle sarcomere. On the other hand, the inter-filament distance in our experiments is not very different from the next-neighbour inter-filament distance between actin filaments in the hexagonal arrangement of thin filaments that surround each thick filament in the sarcomere. In contrast to the inter-head cooperativity involving binding each of the HMM heads between two filaments, we did not study cooperativity of the double-head HMM binding to a filament (similar to in vitro motility or laser-trapping), due to the uncertainty to recognize the binding of specific HMM molecules in subsequent HS-AFM frames (Matusovsky et al., 2021).

### Cooperativity between myosin motors that involves activation of the thin filament

Studies have shown that myosin binding to actin is required for full activation of the thin filament (McKillop and Geeves 1993, Smith & Geeves, 2003; Desai et al., 2015). When myosin binds to actin, it may directly affect the regulatory system by changing the conformation of Tm, such that other myosin heads can attach to thin filaments (Geeves & Holmes, 1999; Gordon, 2000; Smith et al., 2003). According to this model, with the transition from weak to strong actin–myosin binding, the myosin heads transfer Tm to an open state, making neighboring myosin binding sites on actin available for myosin binding. Our data are consistent with this model, as we observed that the binding of one motor to the activated thin filament (pCa 4.5) has changes the attachment kinetics of neighboring motors compared to non-activated thin filaments (pCa 9.0) in the blocked state or the bare actin filaments. Most specifically, when one motor is bound to the activated thin filament at the pCa 4.5, it moves the thin filament from the closed to the open state, which allows for further motor binding at nearby sites.

In a previous study, we showed that the interaction of HMM with cTFs caused a change in the thin filament conformation, both in the absence and presence of Ca^2+^, and in the absence and presence of different concentrations of ATP (Matusovsky et al., 2019).Our new data strengthen those findings and corroborate the idea that cooperativity of myosin heads in striated muscles is defined by thin filaments and their state of activation. We evaluated whether one head in a HMM molecule could activate the thin filament in the presence of ATP at low or high Ca^2+^ concentrations. Under non-activating conditions (presence of ATP, pCa 9.0) when cTFs were in the blocked state, myosin heads were able to bind to cTFs but not able to switch the filaments from the blocked to the closed state, showing a similar decrease in the binding probability and fractional occupancy (Fig. 4e) when compared to the F-actin-HMM complex (Fig.3). Thus, binding of the two myosin heads are required for the transition of a thin filament from the blocked to the close state (Fig. 4). However, the situation is changed if myosin heads bind to cTFs under activation conditions (presence of ATP and pCa 4.5), showing an increase in the probability of binding and the relative number of motors attached to thin filaments, as a result of a first myosin binding (Figs 3 and 4g, Movies S3, S4). These results suggest that one head (upper or lower heads of a given HMM molecule bound between two filaments) is able to switch a thin filament from the closed to the open state.

In addition to the cooperative phenomena considered above, our results also demonstrate higher affinity of myosin heads to the thin filaments in comparison to the actin filaments. This follows from the higher average number of the HMM heads bound (2.32 ± 0.06) to cTFs (Fig. 4h) than to the non-regulated actin filaments (1.27 ± 0.07; Fig. 3f) and the slower decline in the total number of available heads in the former case. These findings are broadly consistent with previous observations that both tension and the average number of attached cross-bridges was increased in actin-reconstituted skinned muscle fibres after furhter reconstitution with thin filament regulatory proteins (Fujita et al, 2002).

To summarize, our data suggest that cooperativity between neighboring myosin molecule along a filament is primarily defined by the state of thin filament activation. In contrast, we find no evidence for cooperative effects attributed to allosteric changes along pure actin filaments.

## Materials and Methods

### Proteins

Native thin filaments were purified from rabbit right and left ventricular cardiac muscle that had been glycerinated and actin was purified from acetone powder of rabbit skeletal muscle (Sigma-Aldrich, USA), following a protocol previously used in our laboratory (Matusovsky et al., 2019). The double-headed skeletal myosin II was purified from rabbit psoas muscle and HMM fragments were prepared by proteolysis of the myosin with α-chymotrypsin (Oakville, Ontario, Canada) as previously described (Cheng et al., 2019). Prior of the HS-AFM experiments, HMM, F-actin and thin filaments were tested for their functionality using *in-vitro* motility and Mg^2+^-ATPase activity assays, as previously described (Matusovsky et al., 2019).

### The lipid bilayer template surface and experimental design

The lipid composition for HS-AFM imaging contained 1,2-Dipalmitoyl-sn-glycero-3-phosphocholine (DPPC, Avanti Polar Lipids), 1,2-Dipalmitoyl-3-trimethylammonium-propane (DPTAP, Avanti Polar Lipids) and 1,2-dipalmitoyl-sn-glycero-3-phosphoethanolamine-N-(cap biotinyl) (biotin-cap-DPPE, Avanti Polar Lipids). DPPC: DPTAP: biotin-cap-DPPE were mixed in a weight ratio of 89:10:1. The preparation of lipid vesicles and deposition on mica substrate to form a mica-supported lipid bilayer surface (mica-SLB) has been previously described (Matusovsky et al., 2021).

The mica-SLB surface was rinsed with the buffer A, containing 25 mM KCl, 2 mM MgCl2, 0.25 mM EGTA, 1.25 mM Imidazole-HCl, 0.5 mM DTT, (pH 7.0). Subsequently, 2.8 µl of either 7 µM non-regulated actin filaments or 1.0 µM regulated cTFs diluted in the buffer A were deposited on the mica-SLB surface and incubated for 10 minutes in a wet chamber. At these conditions, many filaments were attached to the surface in close proximity to each other. The distance distributions between two parallel non-regulated actin filaments and regulated cTFs are shown in Fig.S2. The observed distances were enough for binding the HMM heads between two parallel filaments, allowing counting of the exact number of HMM molecules at the given time of the experiment. We studied cooperativity of binding of the myosin heads in F-actin-HMM or cTF-HMM complexes using the following experimental conditions: i) nucleotide-free (NF) state; ii) presence of ATP analogs (ATP-γ -S); iii) presence of ATP and Ca^2+^.

### HS-AFM imaging of F-actin-HMM complex

After rinsing unbound actin filaments with buffer A, 3.0 µl of 8 nM HMM diluted in buffer A was placed on top of non-regulated actin filaments on the mica-SLB surface, and incubated for an additional 3 minutes. The F-actin-HMM complex in nucleotide-free (NF) conditions was rinsed by 10 μl of buffer A, containing either NPE-caged ATP, non-caged ATP (0.5, 2 or 10 µM) or 0.5 µM non-hydrolyzable ATP-γ-S. NPE-caged ATP (adenosine 5’-triphosphate, P3- (1-(2-nitrophenyl ethyl) ester) (Invitrogen) dissolved in attachment buffer was photolyzed in the AFM chamber using an UV light source at 340 nm. A delay of ∼5-10 seconds was found after activation of caged ATP, likely because caged ATP molecules were in solution and required this time to attach to and get hydrolyzed in the motor domain of HMM. To ensure nucleotide free conditions 1 U/µl of apyrase was added to the solution. Further, 1 U/ml of hexokinase and 10 mM glucose were added to the ADP solutions to remove contaminating ATP. The F-actin-HMM complex was formed on the mica-SLB surface in the buffer A with low (pCa 9.0) or high (pCa 4.5) Ca^2+^ concentrations to ensure similar experimental conditions as for cTFs-HMM complex.

### HS-AFM imaging of cTFs-HMM complex

The procedure for imaging the cTFs-HMM complex was similar to that explained above for non-regulated actin filaments. Imaging of the cTFs-HMM complex at low Ca^2+^ (pCa 9.0) or high Ca^2+^ (pCa 4.5) concentrations, using skeletal muscle HMM was performed in the following way: 2-20 μL of TFs (1 μM) in the buffer A (relaxing conditions, absence of Ca^2+^) were placed on a mica-SLB surface for 10 min in the wet chamber and unbound cTFs were removed by exchanging for the buffer B, containing low or high Ca^2+^ concentrations. Then, 3.0 μL of skeletal muscle HMM (8 nM) in buffer A was placed on top of the mica-SLB surface with bound cTFs for 10 min in the wet chamber. Unbound HMM was washed out by buffer A followed by washing several times with appropriate buffer B with low or high Ca^2+^ concentrations containing 0.5-2 μM of caged or non-caged ATP as desired.

### Probability of binding, fractional occupancies and cooperativity analysis

Probability of the HMM heads binding to the non-regulated or regulated actin filaments was calculated by binomial distribution evaluated in HS-AFM experiments. This assumes that the binding situation in each frame is treated as an independent event because each myosin head is assumed to undergo independent cycling (possibly several times per frame). The bound and unbound events (0 – no binding, 1 – binding) were visualized directly to compute probabilities for the binding-unbinding process as a ratio of the binding events to the total number of events in each independent frame. Our experimental design allows to visualize 4-6 HMM molecules (or 8-12 individual heads) bound between two filaments. Two typical scanning views and rates were used: 150 *×* 75 nm^2^(80 *×* 40 pixels^2^) at the 6.7-10 frame per second (fps) and 200 *×* 200 nm^2^ (120 *×* 120 pixels^2^) at the 2 fps.

Fractional Occupancy (θ) is the ratio of the actin-binding sites occupied by HMM heads to the total number of the actin binding sites experimentally observed in the given time of the experiment and calculated from:

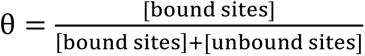

Cooperative probability which is related to the probability of binding was calculated from:

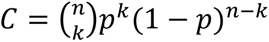

where C denotes cooperative probability of binding, n = number of total events or subsequent frames of the experiment; k = number of binding events in the experiment, p = probability of binding, 1-p = probability of unbinding and 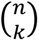 represents the combination of the total and binding events expressed as: 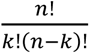.

The events for each myosin head calculated from the reference frame, *i.e.* a moment when the head was bound to the filament until the end of image acquisition. The total events included both binding events (head was bound to actin filament) and unbinding events (head was unbind from actin filament).

### Analysis of the myosin displacements

To analyze the HMM displacement, each of the HMM heads bound between two parallel non-regulated or regulated actin filaments were tracked individually in successive HS-AFM frames. The tracked parameters included the height of the HMM head used for subsequent determination of the center of mass (COM) in each myosin head. The height of the HMM heads was determined in semi-automatic mode using the *x*, *y*, and *z* data of the HS-AFM frames in Kodec software (v. 4.4.7.39) (Ngo et al., 2015). The *x* and *y* data correspond to the lateral coordinates, while the observed *z* values correspond to the highest point in the center of the HMM heads. To obtain the *z* values for the highest point in HMM head(s) the image was automatically searched within a 5 *×* 5 pixels area. Next, the obtained height values and *x*, *y* positions within the 5 *×* 5 pixels area were used to automatically calculate the COM. To obtain the accurate COM values, the height of the surface outside of the actin-HMM position was subtracted from the average COM of the HMM heads. The displacement size was calculated as a difference in the COM position of HMM head in the reference frame and the next frame, in successive HS-AFM frames. The forward and backward displacements were calculated for each myosin head, plotted and fitted by sum of two Gaussians. The displacement size of the upper and lower HMM heads did not differ between each other, although the frequency and binding events were not correlated between two heads within one HMM molecule. The displacement size was also not affected by the range of the ATP concentrations used in our experiments (0.2 μM, 0.5 μM, 2 μM, 10 μM), thus we averaged the data with the sampling rate of 589 events for the non-regulated actin filaments and 911 events for the regulated cTFs.

### Cross-sectional analysis

Cross-sectional analysis was performed by Kodec software (Ngo et al., 2015) to calculate the distance between two filaments (Fig. S2).

### HS-AFM system and cantilevers

The experiments were performed on a tapping-mode HS-AFM system (RIBM) (Ando et al., 2001), equipped with an additional UV laser. Olympus cantilevers BL-AC10DS-A2 with the following parameters were used: spring constant 0.08-0.15 N/m; quality factor in water ∼1.4-1.6; resonance frequency in water 0.6-1.2 MHz. The additional carbon probe tip was fabricated on the tip of a cantilever by electron-beam deposition and sharped by plasma etcher, giving a ∼4 nm tip apex. The tip–sample loading force can be modulated and decreased by the free oscillation peak-to-peak amplitude (A_0_) of the cantilever set to ∼2.0 nm and the amplitude set point adjusted to more than 0.9 A_0_.

### Data analysis and processing of HS-AFM images

To remove spike noise and to make the *xy*-plane flat, the HS-AFM images were processed with low-pass filtering using Kodec software (4.4.7.39). The COM and cross-correlation analyses were performed in Kodec software. Fittings of equations to the observed data were performed in GraphPad Prism software (v.9.3.0). Values are reported as mean ± Standard Deviation or 95% Confidential Intervals throughout the paper as indicated. Number of n equals to independent experiments. A level of significance of p < 0.05 was used for all analyses.

### Data availability

All data required for evaluation of the conclusions in the paper are present in the main body of the paper and/or in the Supporting Information.

## Acknowledgments

This work was supported by the Natural Science and Engineering Research Council of Canada (to D.E.R). A.M. was supported by the Swedish Research Council (grant number 2019-03456). D.E.R. is a Canada Research Chair in Muscle Biophysics. We thank Dr. Y.-S. Cheng for the HMM preparation.

## Author contributions

O.S.M. and D.E.R. designed research; O.S.M. performed HS-AFM experiments and all authors were involved in analysis and interpretation of the data. O.S.M., A.M., D.E.R. wrote the paper and all authors approved the final version of the manuscript.

## Ethics declarations

### Competing interests

The authors declare no competing interests.

## Supplementary Figures

**Figure S1.**
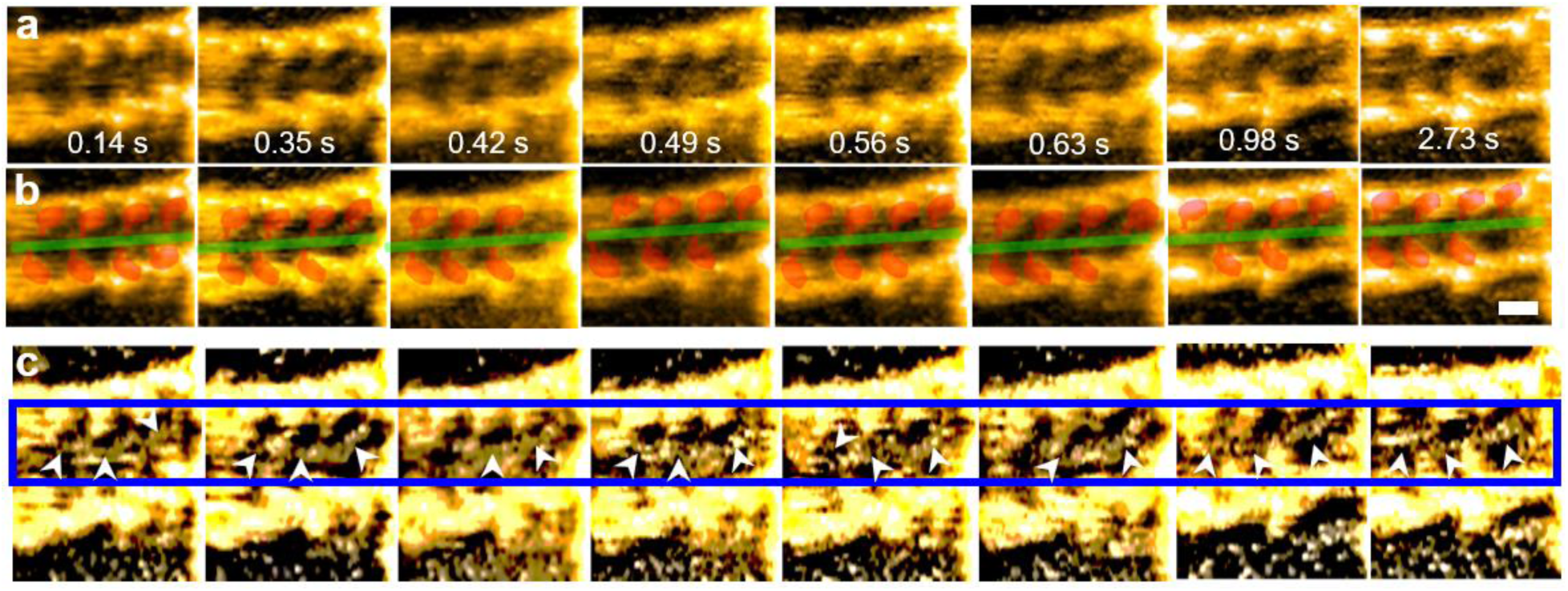
Successive HS-AFM images of F-actin-HMM complex in the presence of 0.2 μM ATP. **(a)** Double-headed heavy meromyosin (HMM) motors bound between two actin filaments in ∼37 nm distance to form a stable structure with up to eight individual myosin heads (four HMM molecules), attached by their S2 regions. (**b**) HMM heads between two actin filaments highlighted by red colors, S2 region of the HMM molecules highlighted by green color. (**c**) high contrast images from (a) panel to highlight the connected S2 regions between HMM molecules. Numbers indicate the time in seconds, the scan rate: 14.4 fps, scale bar: 30 nm.

**Figure S2.**
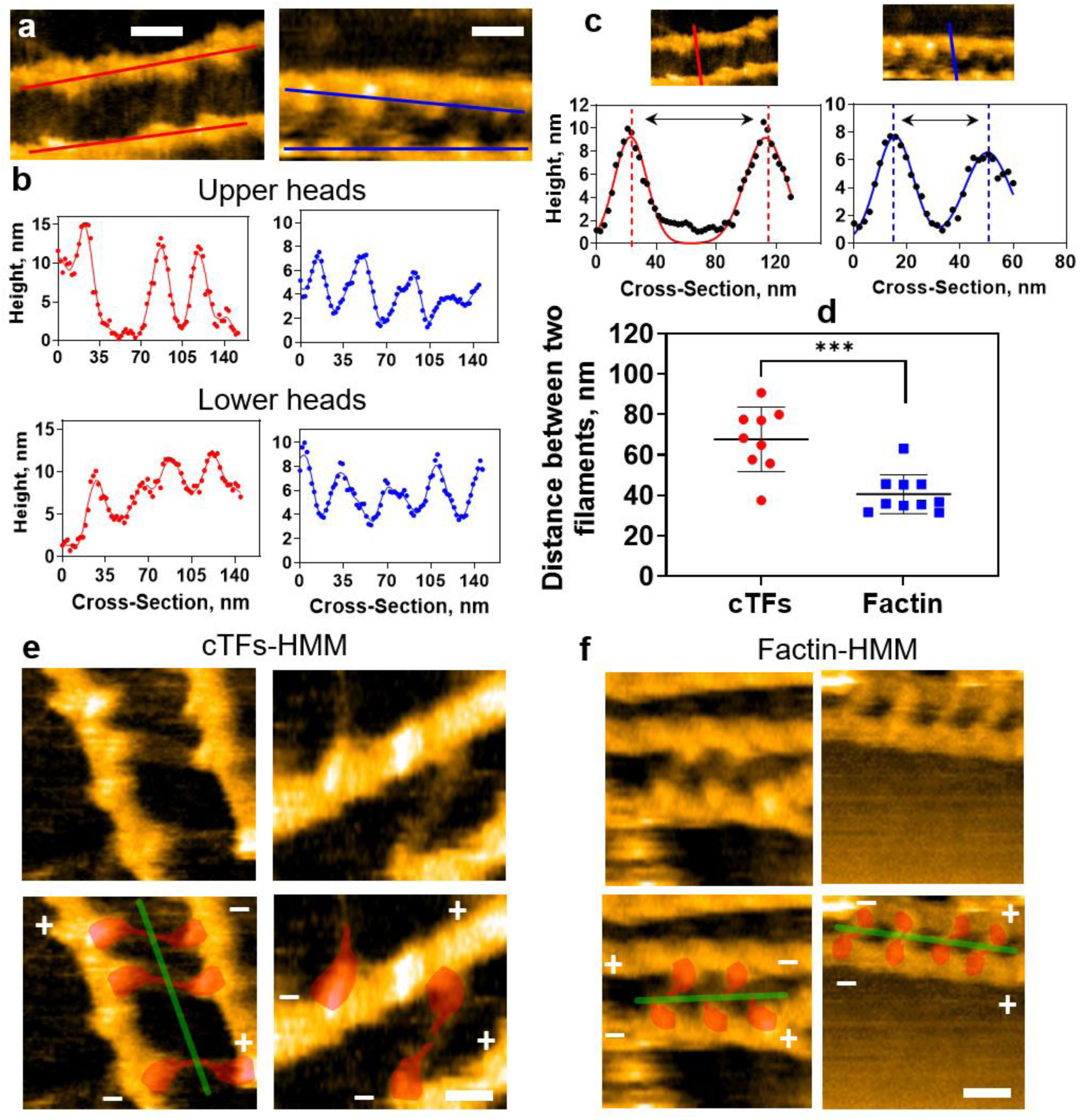
Arrangement of myosin heads between two parallel filaments. (**a-b**) Cross-sections of myosin upper and lower heads bound between two parallel cTFs (left, red profiles) and F-actin (right, blue profiles), scale bars: 30 nm. (**c**) The measured distance between two parallel cTFs (left) and F-actin (right) averaged in (**d**). The difference in distance between non-regulated F-actin and regulated cTFs (p=0.001, unpaired t-test) did not affect the HMM binding and displacement analysis (Figs 1f and 2c). (**e-f**) Two types of HMM binding between parallel cTFs or actin filaments were observed: the upper and lower heads bound towards the similar direction (right HS-AFM images in **e-f**) or the upper and lower heads bound towards the opposite directions (left HS-AFM images in **e-f**). The captured snapshots in left **e** and left **f** panels indicate the rare situation occurred due to head displacement on the filament in the presence of ATP. HMM heads between two actin filaments highlighted by red colors, S2 region of the HMM molecules highlighted by green color. The corresponding polarity of the filaments is shown. Scale bars: 30 nm.

**Figure S3.**
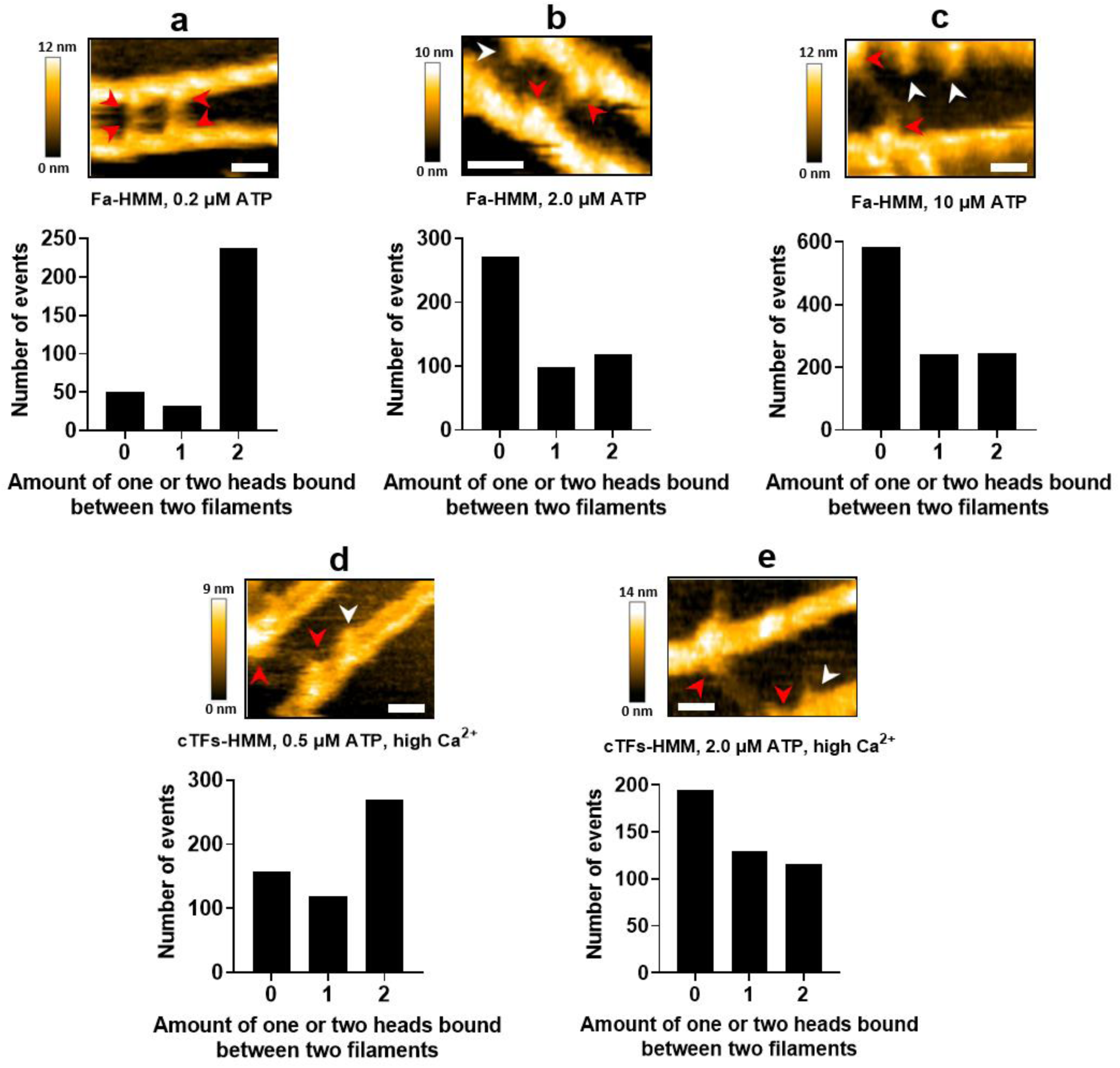
Frequency distribution of one HMM head or two HMM heads bound between two filaments in the presence of ATP. **(a-c)** Distribution of one (1) and two (2) HMM heads in Fa-HMM complex in the presence of 0.2 μM ATP (319 binding events), 2.0 μM ATP (488 binding events) or 10 μM ATP (1072 binding events); **(d-e)** Distribution of one (1) and two (2) HMM heads in cTFs-HMM complex in the presence of 0.5 μM ATP, high Ca^2+^ concentration (545 binding events) and 2.0 μM ATP high Ca^2+^ concentration (440 binding events). The zero indicates the temporarily no bindings of HMM heads along the length of the filaments in analyzed datasets. HMM heads are indicated by white arrows (one head bound) and red arrows (two heads bound). Scale bars: 30 nm; z-scales indicated for each HS-AFM image.

**Figure S4.**
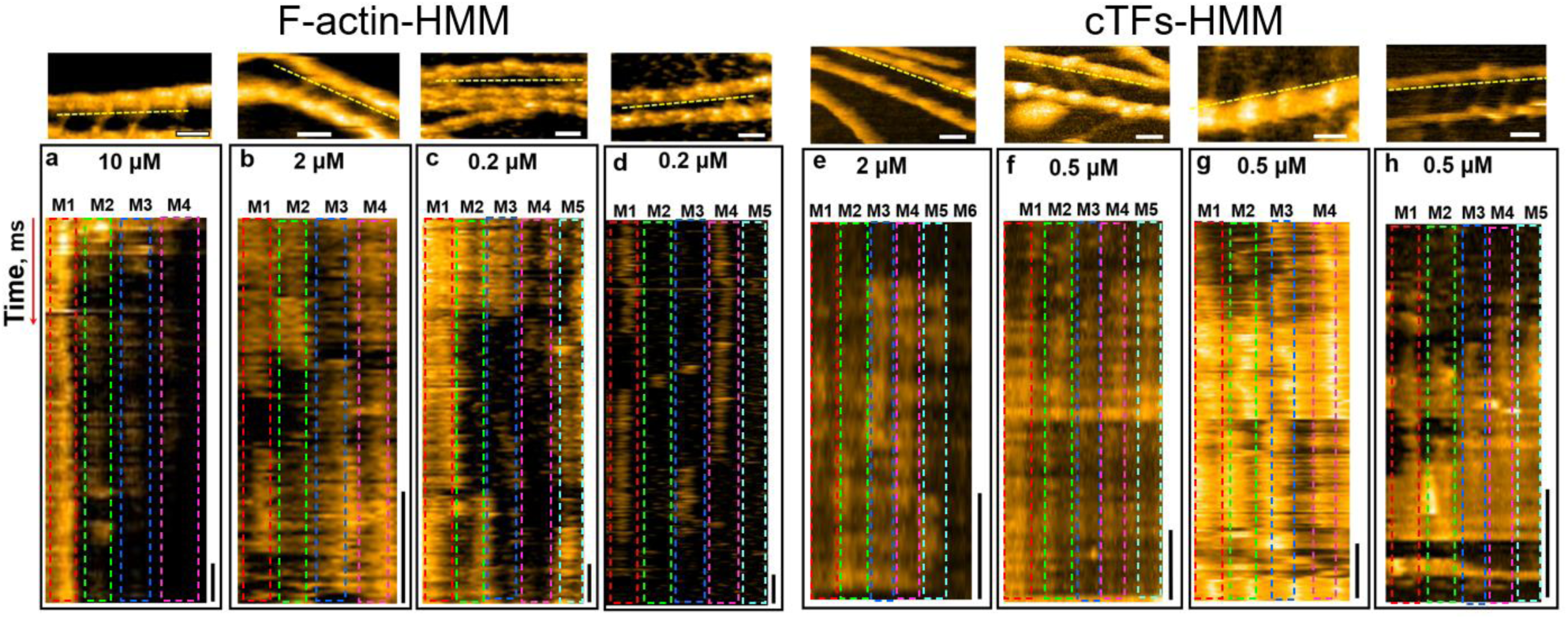
Kymograph images of the F-actin-HMM (a-d) and cTFs-HMM (e-h) complexes at the different ATP concentrations. Scan area and scan rates of the top HS-AFM images for F-actin-HMM complex: (**a**)150 *×* 75 nm^2^, 80 *×* 40 pixels^2^, 10 fps; (**b**) 150 x 90 nm^2^, 80 *×* 48 pixels^2^, 20 fps; (**c**) 200 *×* 120 nm^2^, 80 *×* 40 pixels^2^, 10 fps; (**d**) 200 *×* 120 nm^2^, 80 *×* 40 pixels^2^, 12.5 fps. The horizontal scale bars: 30 nm; the vertical scale bars in kymographs: 20 ms. Scan area and scan rates of the top HS-AFM images for cTFs-HMM complex at the activating conditions: (**e**)120 x 120 nm^2^, 200 *×* 200 pixels^2^, 6.7 fps; (**f**) 120 *×* 120 nm^2^, 200 *×* 200 pixels^2^, 2 fps; (**g**) 150 *×* 75 nm^2^, 80 *×* 40 pixels^2^, 6.7 fps; (**h**) 200 *×* 200 nm^2^, 120 *×* 120 pixels^2^, 2 fps. The horizontal scale bars: 30 nm; the vertical scale bars in kymographs: 50 ms (e-f) and 100 ms (g-h). The dashed yellow line indicated the initial position to create a kymograph image. (**i**) Average dwell-time of myosin heads in F-actin-HMM complex in the presence of 0.5 μM ATP (n=3, 101 events, ∼10 HMM heads) and 2 μM ATP (n=4, 592 events, ∼20 HMM heads). (**j**). Average dwell-time of myosin heads in cTFs-HMM complex in the presence of 0.5 μM ATP (n=3, 825 events, ∼10 HMM heads) and 2 μM ATP (n=3, 498 events, ∼10 HMM heads).

**Figure S5.**
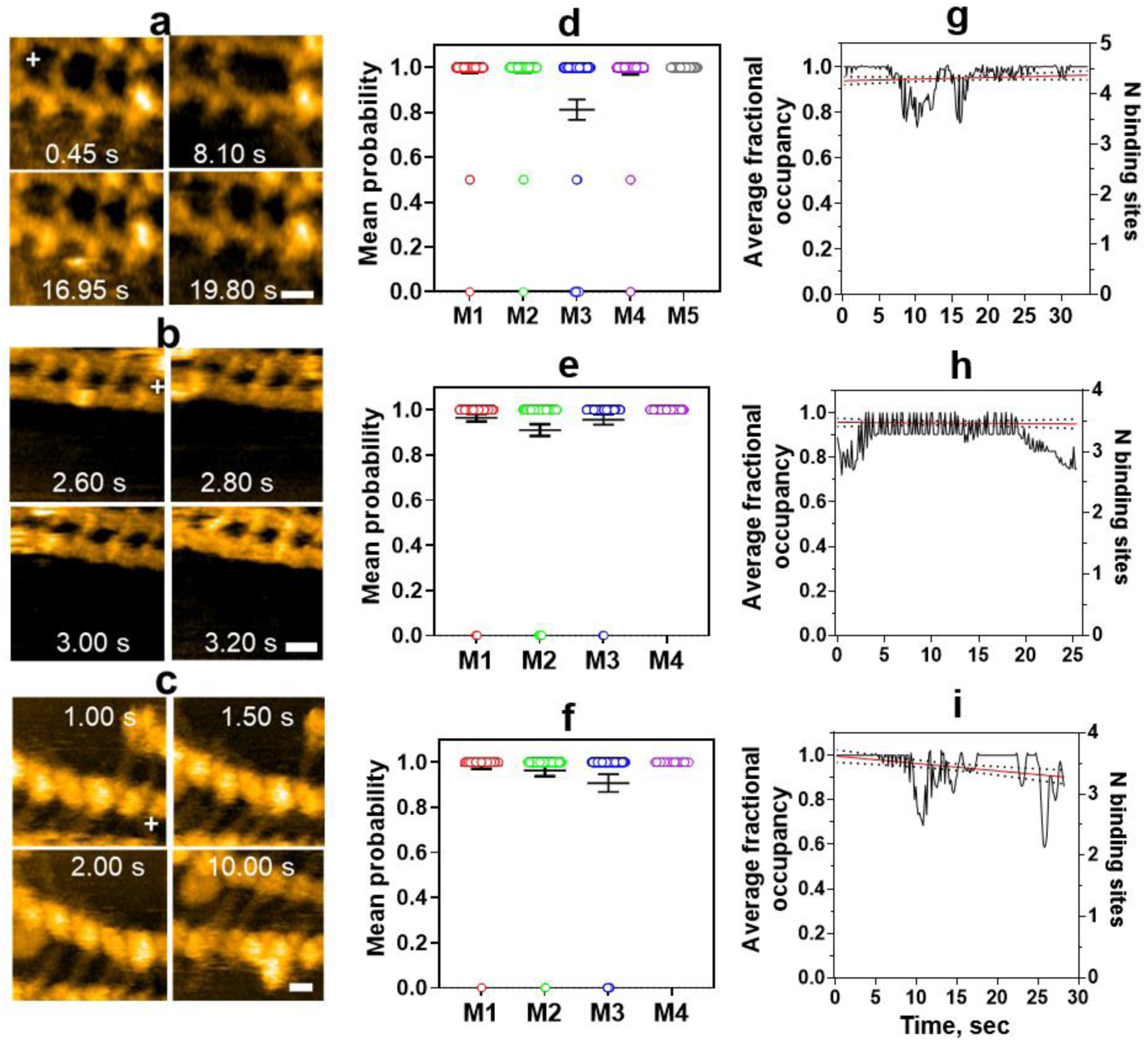
Probability of binding of myosin heads to the non-regulated or regulated filaments in rigor conditions or in the presence of ATP-γ-S. **(a-c)** HS-AFM images of HMM molecules bound between two actin filaments in the presence of ATP-γ-S (**a**), HMM molecules bound between two actin filaments in the absence of Ca^2+^ and ATP (**b**) and HMM molecules bound between two cardiac thin filaments in the absence of Ca^2+^ and ATP (**c**). Scale bars: 30 nm (**d-f**) Probabilities of binding of the HMM heads in the F-actin-HMM complex in the presence of ATP-γ-S (d, 1145 events), in the absence of Ca^2+^ and ATP (e, 1408 events) and in the cTFs-HMM complex in the absence of Ca^2+^ and ATP (f, number of events 663). Individual data points are shown for each HMM molecules (M1-M4) as mean values ± 95% CI, n=3. (**g-i**) Corresponded fractional occupancies of the binding sites by HMM heads for the conditions shown in d-f.

**Figure S6.**
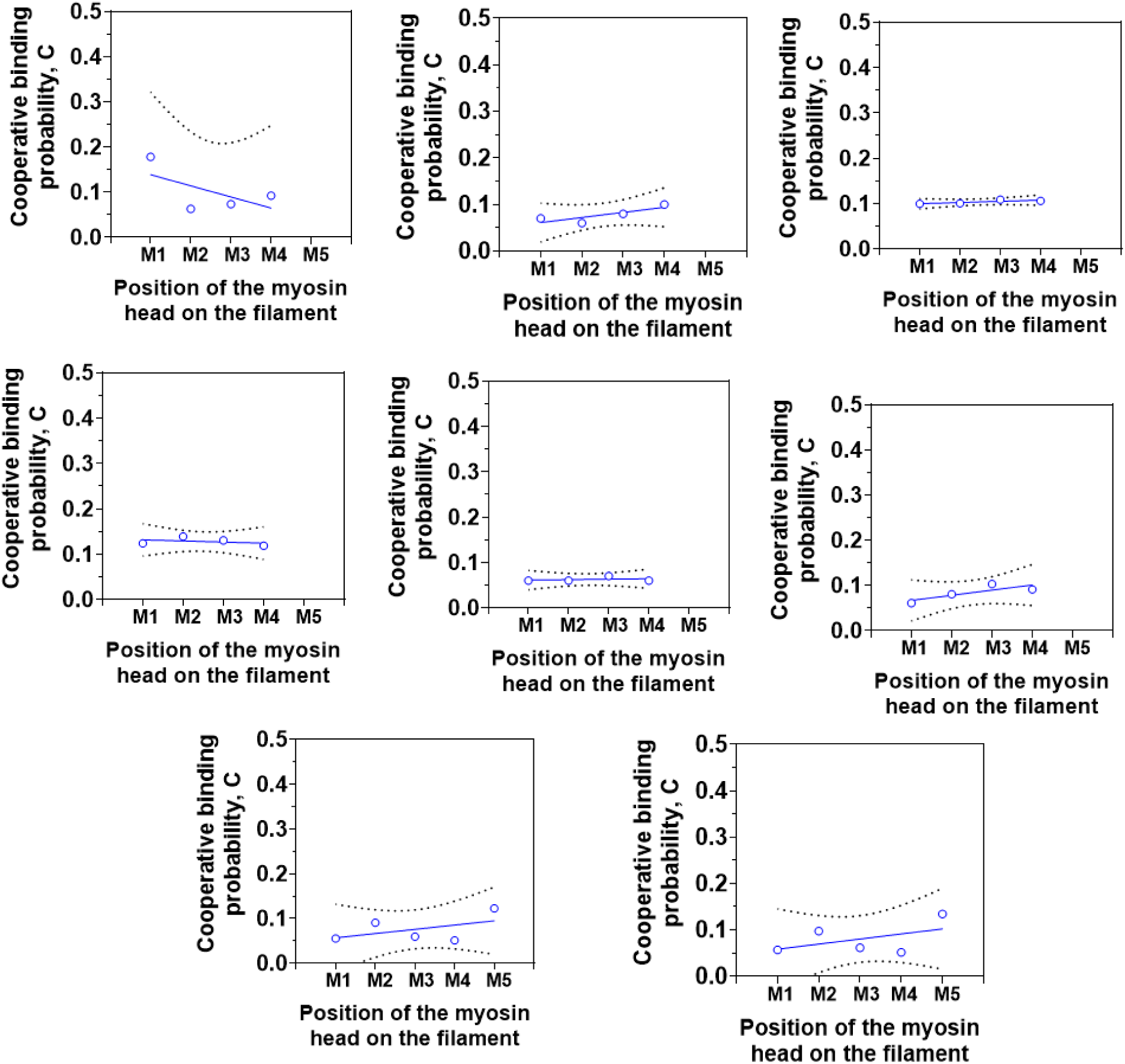
Cooperative probability of the myosin heads bound to the non-regulatory F-actin. The independent experiments showing different patterns of the cooperative probabilities between the one HMM molecule M_n_ and the next molecule M_n+1_ in the F-actin-HMM complex. The linear regression fittings showed in the solid lines and the dashed curves represented 95% CI.

**Figure S7.**
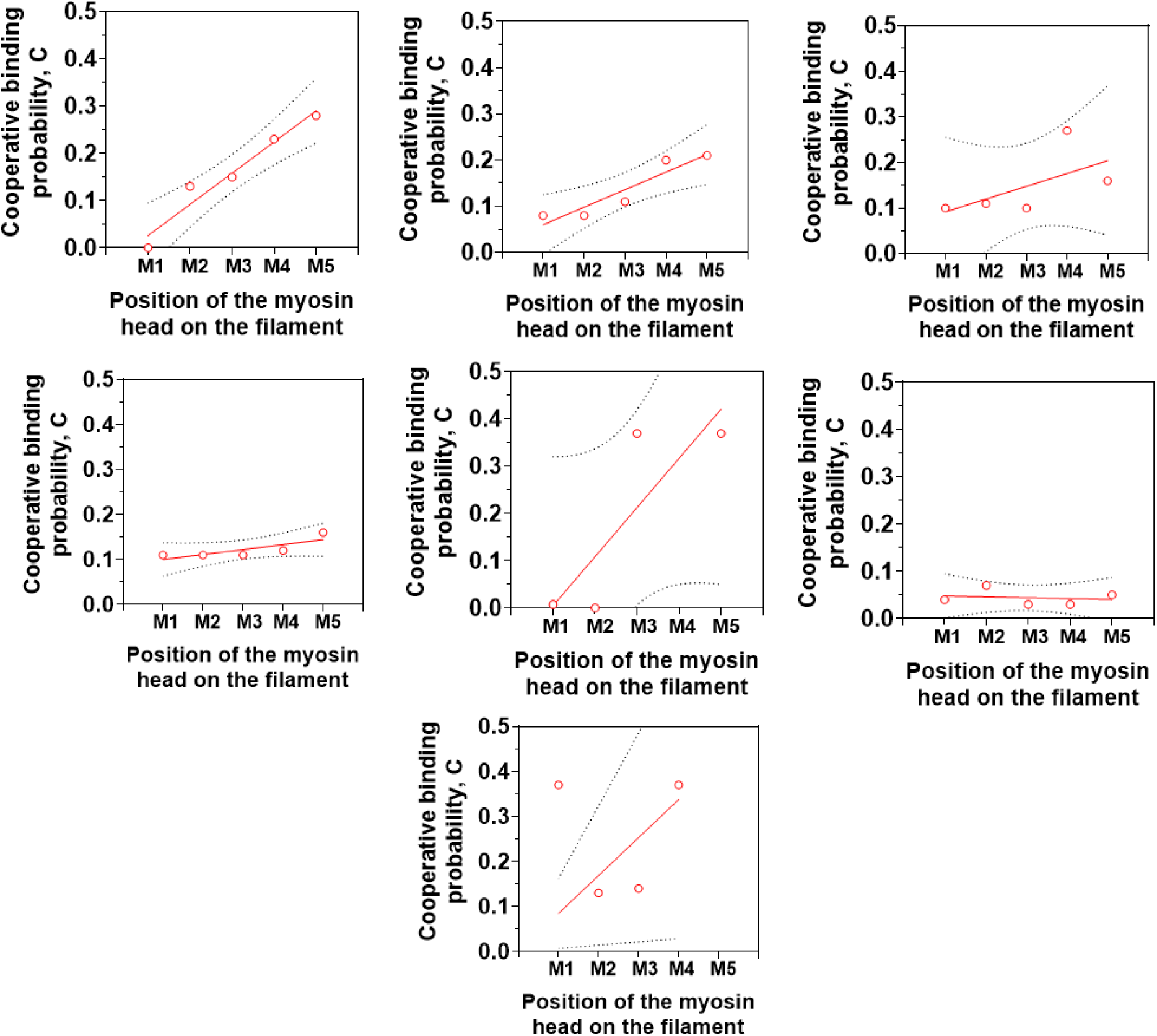
Cooperative probability of the myosin heads bound to the regulatory cardiac TFs. The independent experiments showing different patterns of the cooperative probabilities between the one HMM molecule M_n_ and the next molecule M_n+1_ in the cTFs-HMM complex. The observed differences, most likely related to the diverse population of activated / non-activated segments across the thin filament length that led to different degree of positive cooperativity of myosin bindings. The linear regression fittings showed in the solid lines and the dashed curves represented 95% CI.

To calculate the cooperative probability from observed probability patterns, we applied the following equation 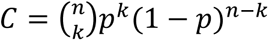

where C denotes cooperative probability of binding, n = number of total events or subsequent frames of the experiment; k = number of binding events in the experiment, p = probability of binding, *i.e.* the ratio between binding events and total events and 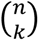 represents the combination of the total and binding events expressed as: 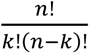.

**Table S1.**
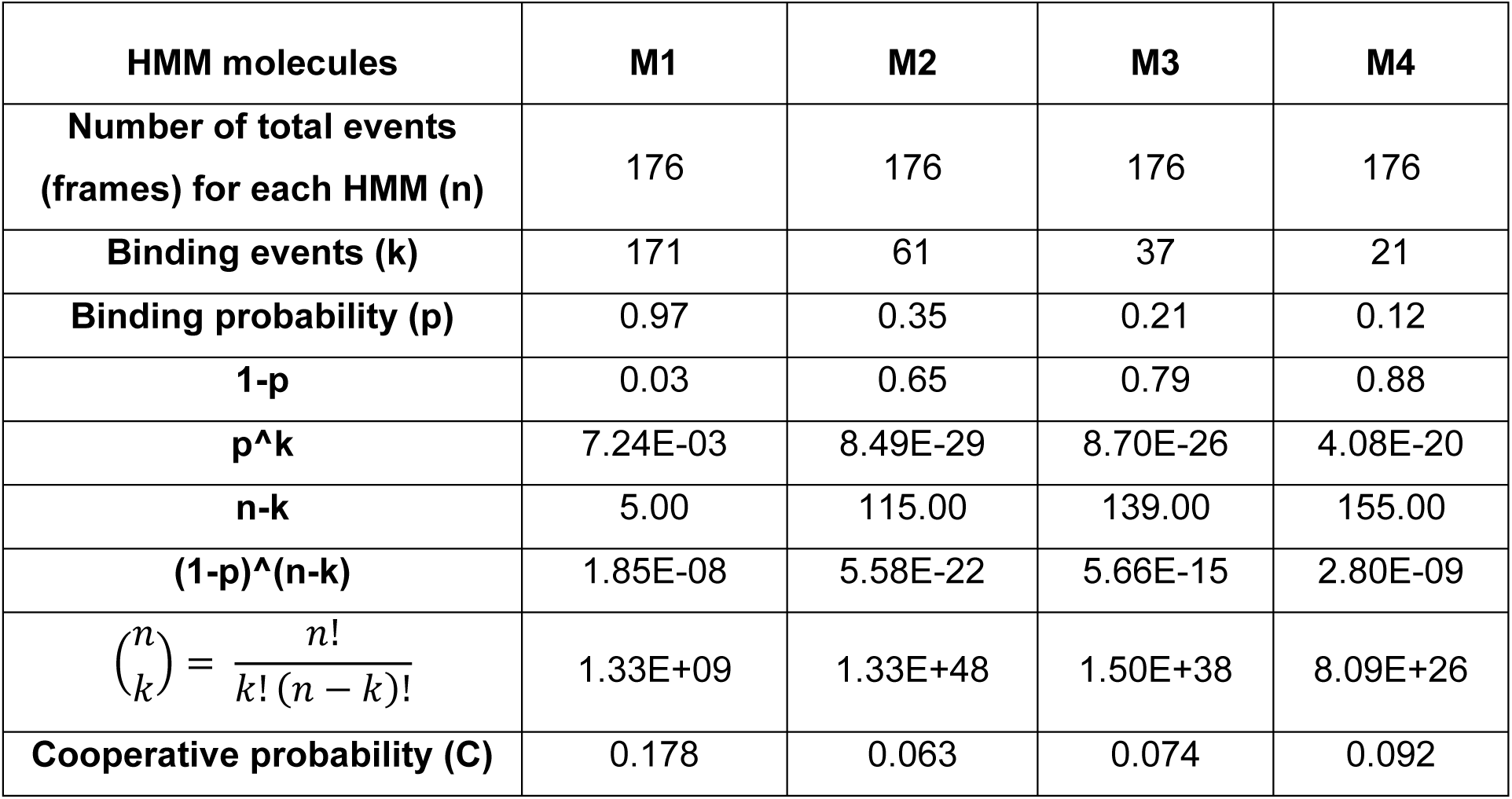
A concrete example of calculation of the cooperativity of binding for 4 neighboring HMM molecules bound to actin filaments in the presence of 10 μM ATP.

## Supplementary Movies

**Movie S1**

Representative HS-AFM movies of the transient binding of skeletal HMM molecules bound between two actin filaments in the presence of 0.2 μM Mg.ATP (top panel). The colored boxes indicated upper and / or lower HMM heads bound to actin filaments. The color code corresponded to the position of the HMM molecule on the actin filaments – 1^st^ molecule (M1): red box; 2^nd^ molecule (M2): green box; 3^rd^ molecule (M3): blue box; 4^th^ molecule (M4): magenta box; 5^th^ molecule (M5): cyan box. The same color code was used in the mean probability binding of the HMM molecules to the actin filaments at the 0.2 μM Mg.ATP (bottom left graph). The average fractional occupation of the actin binding sites in the presence of 0.2 μM Mg.ATP (bottom right graph). Scan area: 150 x 90 nm^2^, 80 x 48 pixels^2^; recording rate 10 fps, playing rate 10 fps (left movie); 200 x 120 nm^2^, 80 x 48 pixels^2^, recording rate 12.5 fps, playing rate 10 fps (central movie); 200 x 120 nm^2^, 80 x 48 pixels^2^, recording rate 10 fps, playing rate 10 fps (right movie). The scale bars are 30 nm.

**Movie S2**

Representative HS-AFM movies of the transient binding of skeletal HMM molecules bound between two actin filaments in the presence of 2 μM Mg.ATP (top panel). The colored boxes indicated upper and / or lower HMM heads bound to actin filaments. The color code corresponded to the position of the HMM molecule on the actin filaments – 1^st^ molecule (M1): red box; 2^nd^ molecule (M2): green box; 3^rd^ molecule (M3): blue box; 4^th^ molecule (M4): magenta box; 5^th^ molecule (M5): cyan box. The same color code was used in the mean probability binding of the HMM molecules to the actin filaments at the 2 μM Mg.ATP (bottom left graph). The average fractional occupation of the actin binding sites in the presence of 2 μM Mg.ATP (bottom right graph). Scan area: 150 x 75 nm^2^, 80 x 40 pixels^2^; recording rate 3.3 fps, playing rate 10 fps; scale bar is 30 nm (left movie); 100 x 60 nm^2^, 80 x 48 pixels^2^; recording rate 6.7 fps, playing rate 10 fps, scale bar is 20 nm (central movie); 100 x 60 nm^2^, 80 x 40 pixels^2^; recording rate 6.7 fps, playing rate 10 fps, scale bar is 20 nm (right movie).

**Movie S3**

Representative HS-AFM movies of the transient binding of skeletal HMM molecules bound between two actin filaments in the presence of 10 μM Mg.ATP (top panel). The colored boxes indicated upper and / or lower HMM heads bound to actin filaments. The color code corresponded to the position of the HMM molecule on the actin filaments – 1^st^ molecule (M1): red box; 2^nd^ molecule (M2): green box; 3^rd^ molecule (M3): blue box; 4^th^ molecule (M4): magenta box; 5^th^ molecule (M5): cyan box. The same color code was used in the mean probability binding of the HMM molecules to the actin filaments at the 10 μM Mg.ATP (bottom left graph). The average fractional occupation of the actin binding sites in the presence of 10 μM Mg.ATP (bottom right graph). Scan area: 150 x 75 nm^2^, 80 x 40 pixels^2^; recording rate 10 fps, playing rate 10 fps (left movie); 150 x 75 nm^2^, 80 x 40 pixels^2^, recording rate 6.7 fps, playing rate 10 fps (central movie); 150 x 75 nm^2^, 80 x 40 pixels^2^, recording rate 10 fps, playing rate 10 fps (right movie). The scale bars are 30 nm.

**Movie S4**

Representative HS-AFM movies of the transient binding of skeletal HMM molecules bound between two cTFs in the presence of 0.5 or 2 μM Mg.ATP (top panel). The colored boxes indicated upper and / or lower HMM heads bound to cTFs. The color code corresponded to the position of the HMM molecule on the thin filaments – 1^st^ molecule (M1): red box; 2^nd^ molecule (M2): green box; 3^rd^ molecule (M3): blue box; 4^th^ molecule (M4): magenta box; 5^th^ molecule (M5): cyan box. The same color code was used in the mean probability binding of the HMM molecules to the cTFs at the 0.5 μM or 2 μM Mg.ATP (middle panel). The average fractional occupations of the binding sites in the presence of 0.5 μM or 2 μM Mg.ATP (bottom panel). Scan area: 200 x 200 nm^2^, 120 x 120 pixels^2^; recording rate 2 fps, playing rate 5 fps (left movie); Scan area: 200 x 200 nm^2^, 120 x 120 pixels^2^; recording rate 2 fps, playing rate 5 fps (central movie); 150 x 75 nm^2^, 80 x 40 pixels^2^, recording rate 6.7 fps, playing rate 10 fps (top right movie); 150 x 75 nm^2^, 80 x 40 pixels^2^, recording rate 6.7 fps, playing rate 10 fps (bottom right movie). The scale bars are 30 nm.

## Notes

### Competing Interest Statement

The authors have declared no competing interest.

